# Lysine deserts prevent adventitious ubiquitylation of ubiquitin-proteasome components

**DOI:** 10.1101/2022.12.08.519562

**Authors:** Caroline Kampmeyer, Martin Grønbæk-Thygesen, Nicole Oelerich, Michael H. Tatham, Matteo Cagiada, Kresten Lindorff-Larsen, Wouter Boomsma, Kay Hofmann, Rasmus Hartmann-Petersen

**Author notes:** Corresponding authors: R.H.-P., K.H., W.B. These authors contributed equally to this work (Caroline Kampmeyer, Martin Grønbæk-Thygesen).

## Abstract

In terms of its relative frequency, lysine is a common amino acid in the human proteome. However, by bioinformatics we find hundreds of proteins that contain long and evolutionarily conserved stretches completely devoid of lysine residues. These so-called lysine deserts show a high prevalence in intrinsically disordered proteins with known or predicted functions within the ubiquitin-proteasome system (UPS), including many E3 ubiquitin-protein ligases and UBL domain proteasome substrate shuttles, such as BAG6, RAD23A, UBQLN1 and UBQLN2. We show that introduction of lysine residues into the deserts leads to a striking increase in ubiquitylation of some of these proteins. In case of BAG6, we show that ubiquitylation is catalyzed by the E3 RNF126, while RAD23A is ubiquitylated by E6AP. Despite the elevated ubiquitylation, mutant RAD23A appears stable, but displays a partial loss of function phenotype in fission yeast. In case of UBQLN1 and BAG6, introducing lysine leads to a reduced abundance due to proteasomal degradation of the proteins. For UBQLN1 we show that arginine residues within the lysine depleted region are critical for its ability to form cytosolic inclusions. We propose that selective pressure to avoid lysine residues may be a common evolutionary mechanism to prevent unwarranted ubiquitylation and/or perhaps other lysine post-translational modifications. This may be particularly relevant for UPS components as they closely and frequently encounter the ubiquitylation machinery and are thus more susceptible to non-specific ubiquitylation.

## Introduction

Ubiquitylation is a post-translational modification (PTM) where a ubiquitin moiety is covalently linked to a target protein^1^. The most common ubiquitin attachment site is the ε-amino group of a lysine residue within the target. The process is catalyzed by three distinct enzymatic activities. First, a thioester bond is created between the C-terminal carboxyl of ubiquitin and a cysteine residue in the ubiquitin-activating enzyme (E1), driven by ATP hydrolysis. Next, the activated ubiquitin is transferred to a ubiquitin-conjugating enzyme (E2). Finally, a substrate-specific ubiquitin ligase (E3) interacts with the ubiquitin-loaded E2 and the target protein and catalyzes the ligation of ubiquitin to a lysine residue in the target protein. Repeated cycles of this ubiquitylation cascade lead to the formation of poly-ubiquitin chains, which target proteins for degradation by the 26S proteasome^2^ or work as a signal in other processes, including DNA repair and endocytosis^3^. For degradation, ubiquitylated target proteins are either directly recognized and bound by the 26S proteasome via its ubiquitin-binding subunits or, alternatively, bound by UBL-UBA shuttle proteins. Through C-terminal ubiquitin-associated (UBA) domains^4^, the UBL-UBA proteins interact with the ubiquitylated targets, while the N-terminal ubiquitin-like (UBL) domains interact with the proteasome^5–7^. The most prominent proteasomal shuttle proteins are yeast Rad23 and Dsk2, and their human orthologues RAD23AB and UBQLN1-4^8^, respectively. Another group of substrate shuttles is the UBL-BAG domain proteins, which include the human BAG1 and BAG6^9^ proteins. Unlike the UBL-UBA shuttles, these do not bind directly to ubiquitin. Instead, they contain a C-terminal Bcl-2 associated athanogene (BAG) domain that links to chaperones, thus bringing the chaperone and its bound substrate within proximity of the proteasome for degradation^10–12^.

Although lysine is a common amino acid (5.6% occurrence in the eukaryotic proteome^13^), a few proteins wholly or largely without lysine residues have been reported, including certain viral proteins^14^ and toxins^15^. In addition, the yeast E3 ubiquitin-protein ligases San1 and Slx5/8 both contain long stretches devoid of lysine, so-called lysine deserts, by which they avoid auto-ubiquitylation^16,17^. By bioinformatics, long lysine depleted regions have also been observed in other E3s^18^. However, it is unknown whether there is a general functional significance of lysine depletion. It has been speculated^14^ that the viral proteins and toxins have also evolved lysine deserts to avoid ubiquitylation and degradation. In case of viral proteins, this would reduce MHC-I-mediated antigen presentation and thus limit their detection by the immune system. In addition, the viral protein lysine deserts might also protect them from conjugation to other lysine-targeting modifications such as ISG15, which inhibits viral infection^19^. For toxins that enter the cytosol through retrograde transport^20^, the lysine deserts may allow them to escape ER-associated degradation (ERAD) as shown for ricin^21^, cholera^22^ and pertussis^23^ toxin, thus resulting in increased potency. Lastly, lysine desert proteins are enriched in actinobacteria and the phages that infect them^46^. This enrichment correlates with the pupylation system found only in this phylum, which, similar to the UPS, involves conjugation to lysine residues as a signal to facilitate degradation^24^.

Using bioinformatics, we identify a wide range of proteins across different species that contain long stretches completely devoid of lysines. Moreover, we analyze lysine deserts in the human proteome and find that they are conserved and pervasive for several proteins with known or predicted roles in the ubiquitin-proteasome system (UPS). Of these, we tested the consequences of artificial lysine introduction for a selected group of proteins (RAD23A, UBQLN1, UBQLN2, BAG6, RNF115 and PSMF1) and found increased ubiquitylation for most of the mutant proteins compared to their wild-type counterparts. In case of BAG6, we show that ubiquitylation is catalyzed by the E3 RNF126, while RAD23A is ubiquitylated by E6AP. Despite the elevated ubiquitylation, mutant RAD23A appears stable but displays a partial loss of function phenotype in fission yeast. In case of UBQLN1 and BAG6, introducing lysine leads to a reduced abundance due to proteasomal degradation of the proteins.

## Results

### A bioinformatics screen for lysine desert proteins

Although lysine desert proteins have been observed before^16–18^ it remains unknown how widespread they are throughout proteomes. To this end, we used a bioinformatics approach to search the entire human proteome for proteins containing stretches without lysine residues and then sorted the proteins based on the length of the lysine depleted region (the lysine desert) (Supplemental file 1, sheet 1). Since many of the highest scoring proteins would likely include proteins with simple repeated sequences lacking lysine, we next filtered for sequence complexity using Shannon entropy^25^ for the lysine desert regions (Supplemental file 1, sheet 2). Then, we annotated the top 200 hits (based on desert length in number of amino acids) for their subcellular localization as provided in UniProt (Supplemental file 1, sheet 3). Most of the lysine desert proteins were known or predicted to be localized to the plasma membrane (41%) or the extracellular matrix (24%), while the cytosol and nucleus accounted for only 15% and 16%, respectively (Fig 1A). Remarkably, out of the cytosolic and nuclear lysine desert proteins, about one third of the lysine deserts were found in proteins with gene ontology (GO) annotations for biological process and molecular function connected with the ubiquitin-proteasome (UPS) system (Fig. 1A and Supplemental file 1). The abundance of UPS components among the lysine desert proteins is also evident without preselecting the cytosolic and nuclear proteins. Among the top 100 and the top 200 lysine desert proteins, 16% and 12%, respectively, are annotated as UPS components, corresponding to a four- and three-fold enrichment relative to the 4% of the human proteome (Fig. 1A). This selection for UPS components was even more pronounced when we repeated our analyses for the proteomes of *Saccharomyces cerevisiae, Schizosaccharomyces pombe, Arabidopsis thaliana, Caenorhabditis elegans* and *Drosophila melanogaster* (Supplemental file 2). For instance, of the proteins with lysine deserts longer than 200 residues (after Shannon filtering for low-complexity regions) in the yeasts *S. cerevisiae* and *S. pombe*, 7 out of 14 (50%) and 17 out of 35 (48%), respectively, are UPS components (Supplemental file 2). The previously described budding yeast lysine desert proteins Slx5 and San1^16,17^ were the highest scoring hits in *S. cerevisiae* (Supplemental file 2)^26,27^.

**Fig. 1.**
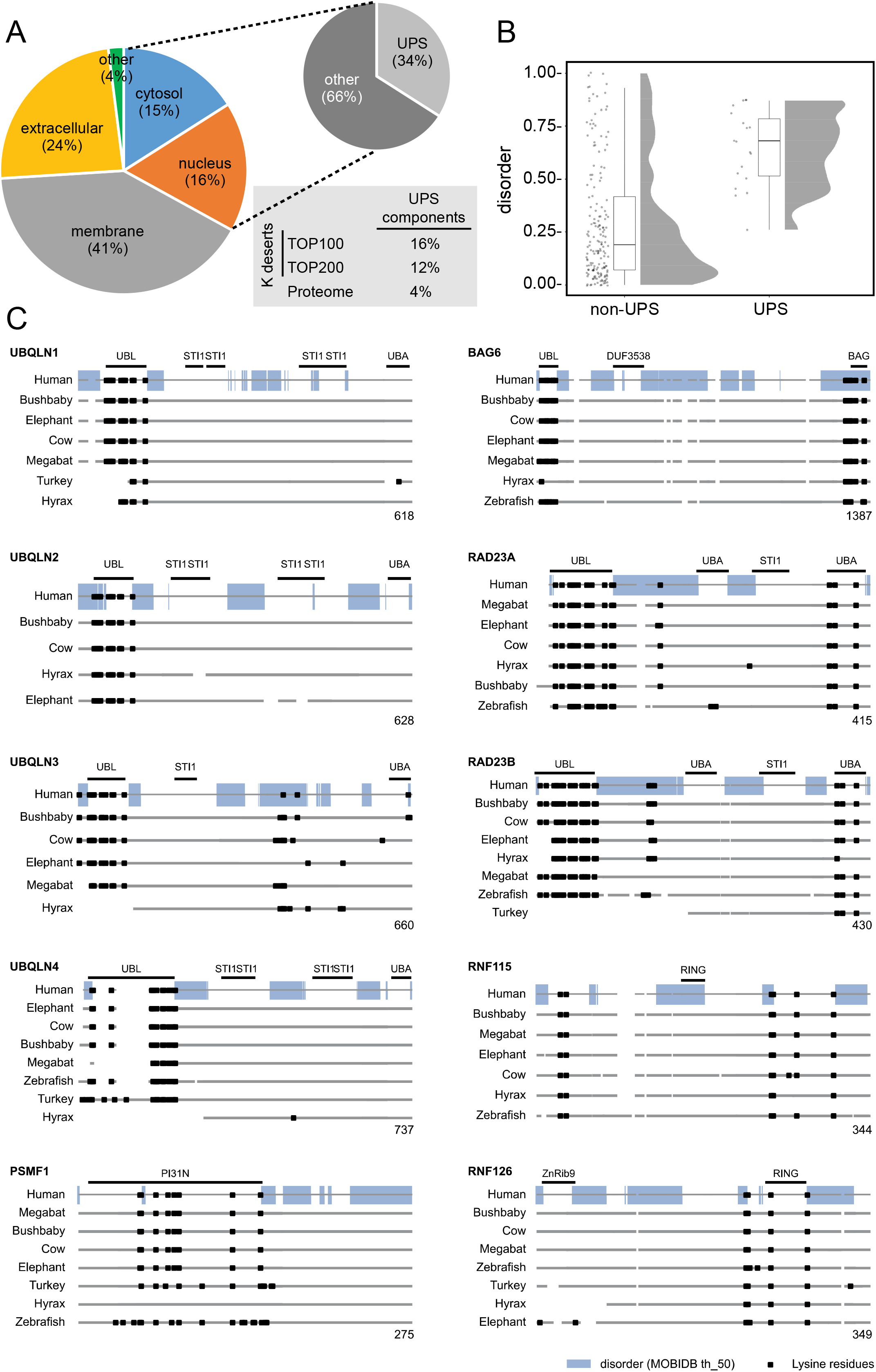
A bioinformatics screen for lysine desert proteins. (A) Pie diagram showing the distribution of the subcellular localization of the top 200 identified human lysine deserts proteins after Shannon entropy filtering. Of the cytosolic and nuclear proteins, gene ontology (GO) classifications link 34% of the proteins to functions within the ubiquitin-proteasome system (UPS). The insert shows that the number of proteins with known or predicted functions in the UPS is also enriched for the top 100 and top 200 lysine desert proteins compared to the entire human proteome. (B) The intrinsic disorder according to the MobiDB database for the top 200 lysine deserts. (C) Conservation of the listed lysine desert proteins in the indicated species. Lysine residues are marked as black squares. The intrinsically disordered regions based on MobiDB in the human proteins are shown as a blue bar. The domain organization according to the SMART database is marked.

To be functionally relevant, a lysine desert likely must be exposed. Accordingly, the lysine deserts in the San1 and Slx5 E3s are localized in intrinsically disordered regions (IDRs)^16,17^. Therefore, we next analyzed the top 200 human lysine desert proteins for intrinsic disorder within the lysine deserts, noting that lysines in general are modestly enriched in IDRs^26,27^. Using the MobiDB database^28^ annotation of intrinsic disorder, the average disorder for proteins in the human proteome was 26%, or 0.26 ± 0.23 (mean ± standard deviation). For the non-UPS components in the top 200, this was not significantly different (0.29 ± 0.28). However, for the UPS components in the top 200 list the average disorder was higher (0.64 ± 0.18) (Fig. 1BC). This shows that the lysine deserts in UPS components often, but not exclusively, overlap with regions of intrinsic disorder.

In agreement with our previous bioinformatics analyses^18^, there are, among the UPS components in the top 200 lysine desert proteins (Table 1), several known or predicted E3s, including AMBRA1, RNF111 and RNF6. We also noted several non-E3 UPS components, including UBQLN1-4 and BAG6. Importantly, since the employed bioinformatics screen selects purely on the length of the lysine desert, deserts located in short proteins are overlooked. Conversely, long lysine deserts in large proteins will be detected but may only cover a smaller fraction of the protein sequence. For instance, the HECD1 E3 contains a 253-residue stretch devoid of lysine residues. However, since HECD1 is 2610 residues long, the desert only accounts for about 10% of the entire protein, but still covers most of the unstructured part of the protein. Accordingly, in the following we analyzed selected UPS lysine desert proteins (RAD23A, UBQLN1, UBQLN2, BAG6, RNF115 and PSMF1) identified broadly within the top 500, but all with lysine deserts covering at least 45% of the entire protein.

**Table 1.** Highest scoring human lysine desert UPS components.

| **Name** | **UniProt ID** | **Desert length (aa)** | **Desert length (%)** | **Localization** | **Function** |
| --- | --- | --- | --- | --- | --- |
| BAG6 | P46379 | 950 | 83.9 | Nucleus | Shuttle |
| AMBRA1 | Q9C0C7 | 693 | 53.4 | Other | E3 co-factor |
| RNF111 | Q6ZNA4 | 663 | 66.7 | Nucleus | E3 |
| RNF6 | Q9Y252 | 527 | 76.9 | Cytosol | E3 |
| UBQLN2 | Q9UHD9 | 521 | 83.5 | Cytosol | Shuttle |
| UBQLN4 | Q9NRR5 | 514 | 85.5 | Cytosol | Shuttle |
| UBQLN1 | Q9UMX0 | 482 | 81.8 | Cytosol | Shuttle |
| RNF12 | Q9NVW2 | 477 | 76.4 | Nucleus | E3 |
| HUWE1 | Q7Z6Z7 | 439 | 10.0 | Cytosol | E3 |
| HECW2 | Q9P2P5 | 393 | 25.0 | Cytosol | E3 |
| RNF38 | Q9H0F5 | 371 | 72.0 | Nucleus | E3 |
| UBR5 | O95071 | 342 | 12.2 | Nucleus | E3 |
| MARCHF7 | Q9H992 | 329 | 46.7 | Cytosol | E3 |
| RNF43 | Q68DV7 | 324 | 41.4 | Nucleus | E3 |
| HDAC6 | Q9UBN7 | 317 | 26.1 | Cytosol | Deacetylase |
| UBQLN3 | Q9H347 | 311 | 47.5 | Cytosol | Shuttle |
| ZSWIM8 | A7E2V4 | 296 | 16.1 | Cytosol | E3 co-factor |
| ANKIB1 | Q9P2G1 | 284 | 26.1 | Cytosol | E3 |
| UBR4 | Q5T4S7 | 275 | 5.3 | Cytosol | E3 |
| HERP1 | Q15011 | 274 | 70.1 | Membrane | UBL protein |
| RNF26 | Q9BY78 | 270 | 62.4 | Cytosol | E3 |
| RNF165 | Q6ZSG1 | 268 | 77.5 | Nucleus | E3 |
| RFWD3 | Q6PCD5 | 258 | 33.3 | Nucleus | E3 |
| HECD1 | Q9ULT8 | 253 | 9.7 | Cytosol | E3 |

### The selected lysine deserts are conserved between species

It is possible that regions with low lysine counts occur through random chance rather than as a result of evolutionary selection. To investigate further whether the regions with lysine depletion are likely to be biologically relevant, we compared the lysine content for selected proteins across a diverse range of orthologues (Fig. 1C). We observe that generally, the lysine deserts are conserved across species, suggesting there was selective pressure for this trait.

To evaluate a potential selection against introducing lysines into lysine deserts, we performed sequence analyses to predict the mutational landscapes of lysine desert proteins by modeling the evolutionary history of their sequences. Specifically, we used GEMME^29^ (Global Epistatic Model for predicting Mutational Effects) and, as an input, multiple sequence alignments of RAD23A, UBQLN1 and BAG6 (with 1040, 623, and 169 entries, respectively) to calculate a score (available in supplemental file 3) for substituting the wild-type residue to all 19 other residue types. A score around zero suggests that a substitution would be acceptable (because it has been observed in this position and context during evolution), whereas more negative scores represent substitutions that the evolutionary model suggest would be detrimental. The evolutionary pressure to keep these regions without lysine was also evident from these analyses (Fig. 2). Thus, in general, these predictions indicate that substitutions to lysine are less tolerated in these proteins (Fig. 2). For RAD23A, UBQLN1 and BAG6, the GEMME scores indicated that substitutions to lysine were more unfavorable than substitutions to arginine (Fig. S1). Orthogonally, we compared the conservation of these three proteins with a set of ten proteins with short lysine desert regions (non-desert proteins). This reveals a small shift toward conservation in the mean value for substitution from any amino acid to lysine within lysine desert proteins compared to non-desert proteins (Fig. S2). An example heatmap of a non-desert UPS protein did not reveal substitutions to lysine as particularly detrimental (Fig. S3). Importantly, this shows that substitutions to lysine is not generally disfavored in eukaryotic proteins, whereas the observed selection against substitutions to bulky hydrophobic residues (W, V and Y) and proline (Fig. 2 and Fig. S3) is a common feature for most structured proteins^30^. However, we also note, in particular in RAD23A and in UBQLN1, that cysteine is unfavorable at most positions (Fig. 2). Indeed, cysteine is rarely found in RAD23A and UBQLN1 orthologues, which we speculate is because cysteine is generally rare in IDRs^31^. Conversely, we note that lysine is common in IDRs^26,27^.

**Fig. 2.**
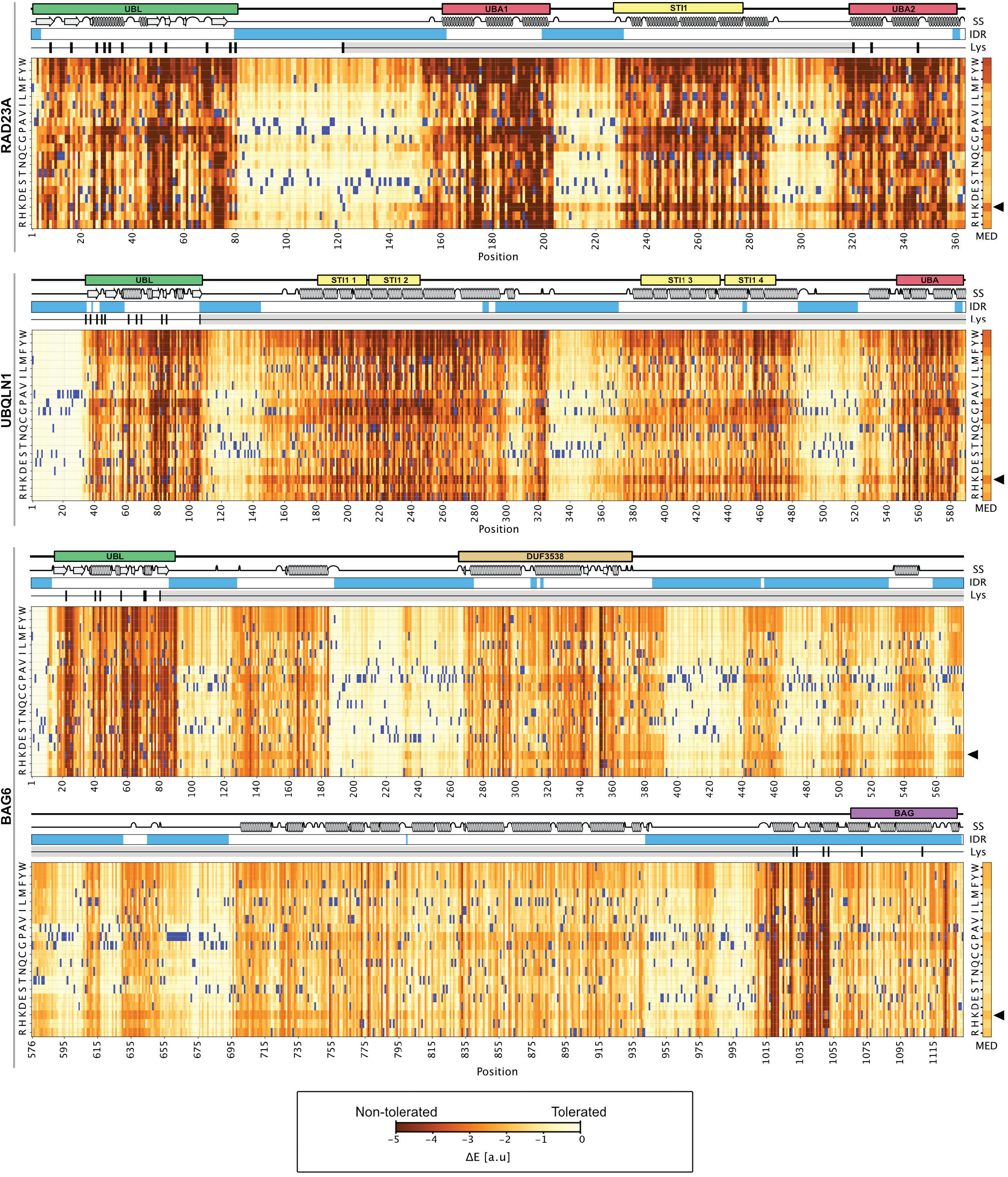
Evolutionary conservation of the lysine deserts. Evolutionary conservation analysis using multiple sequence alignments of human RAD23A, UBQLN1 and BAG6 presented as heat maps. Scores (ΔE) close to zero (white and light yellow colors) indicate that a given amino acid substitution is compatible with the alignments, while negative scores (red and dark yellow colors) indicate that the substitution is incompatible with the alignments and therefore likely detrimental to the protein structure and/or function. The wild-type amino acid residue at each position is shown in blue. The domain organization, intrinsic disordered regions (IDR) (blue) and structural elements are shown above. The positions of lysine residues (black bars) and the extent of the lysine deserts (grey bars) is indicated. The median score (MED) for substitutions to the indicated amino acid resides across the entire protein is shown to the right. Note that substitutions to lysine (arrowhead) in general appear detrimental.

The human UBL-UBA shuttle RAD23A carries a 198-residue uninterrupted central lysine desert spanning the protein from the N-terminal UBL domain to the C-terminal UBA domain. UBLQN1 and UBQLN2 are even more dramatic, with their entire sequences downstream of the N-terminal UBL domain, corresponding to 482 and 521 residues, respectively, without any lysine. BAG6 is a massive protein of 1132 residues with a central uninterrupted desert of 950 residues without lysine. RNF115 is a 304-residue long RING-type E3 carrying a disordered N-terminal region with a desert of 173 residues. Finally, PSMF1, also known as PI31, is a 20S proteasome inhibitor^32^ in which the entire disordered C-terminal region of 123 residues is a lysine desert. Importantly, in these cases the regions are also depleted for lysine in their human paralogues (RAD23B, UBQLN3 and UBQLN4), despite an otherwise high sequence variability in the disordered regions. Finally, since these lysine deserts are not depleted of arginine (Fig. 3A), the lack of lysine is unlikely to be a consequence of an evolutionary pressure to maintain a specific overall net charge. This implies that the lysine deserts are rather a consequence of avoiding a functionality specific to the lysine side chain, which is undesirable in these regions.

**Fig. 3.**
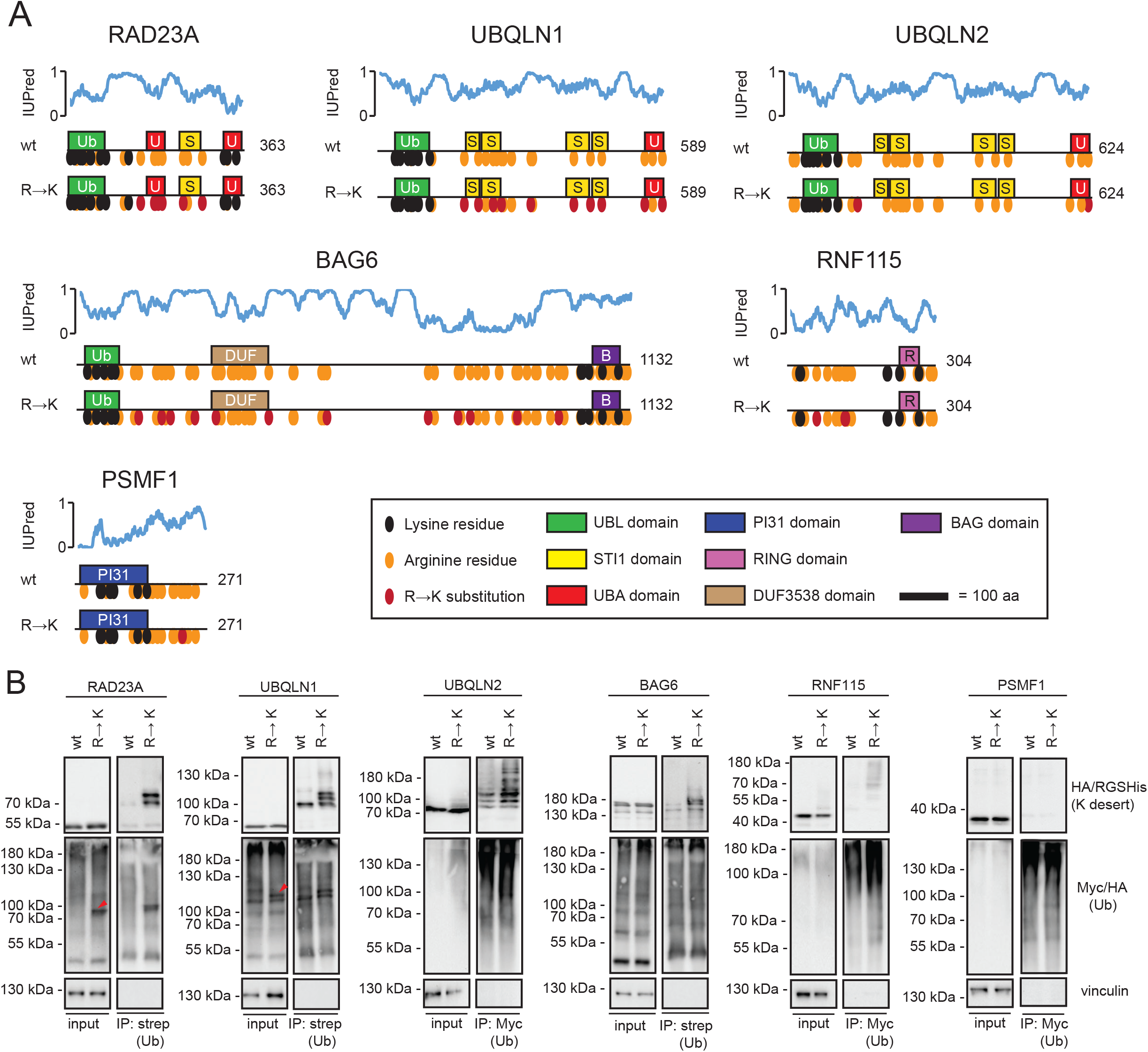
The introduction of lysine residues leads to increased ubiquitylation of UPS components. (A) Schematic illustration of the indicated constructs, showing the positions of domains to scale, endogenous arginine (orange) and lysine (black) residues and introduced R→K substitutions (red). The prediction of intrinsically unstructured regions was performed with the IUPred web server (https://iupred.elte.hu/)^78^. (B) U2OS cells were transiently co-transfected with either HA-strep- or Myc-tagged ubiquitin and the indicated constructs. After 24 h, cells were treated with 10 μM BZ for 16 h and then used for denaturing immunoprecipitation (IP) of either strep or Myc. Protein was visualized by Western blotting and vinculin was used as a loading control. Note that the ubiquitylation of RAD23A and UBQLN1 R→K variants was evident already in the input samples (red arrows).

### Introduction of lysine residues into lysine deserts leads to increased ubiquitylation

One function of lysine residues is to accept covalent ubiquitin modifications. To test if the introduction of lysine residues into lysine deserts leads to altered ubiquitylation of the corresponding mutant protein, we transiently co-transfected human U2OS cells to express tagged ubiquitin (HA-strep- or myc-tagged) and one of six selected proteins, either wild-type or a version where one or multiple arginine residues had been substituted for lysine (R→K) (Fig. 3A). The proteasome inhibitor bortezomib (BZ) was added to the cells 16 h prior to harvest to ensure that ubiquitylated proteins were not degraded. To purify only proteins covalently conjugated to ubiquitin, and not ubiquitin interacting proteins, the cell lysates were first denatured by incubation at 100 °C in SDS. Then, SDS was removed with Triton X-100 prior to precipitation of the tagged ubiquitin. We detected an increased ubiquitylation of the R→K variants of RAD23A, UBQLN1, UBQLN2, BAG6 and RNF115 compared to their wild-type counterparts, but not of PSMF1 (Fig. 3B). Furthermore, in the precipitates of the K variants, the ubiquitylated proteins all had a higher molecular weight, supporting that the bands represent covalent conjugates. These results were specifically dependent on the introduction of lysine, as control experiments, where the arginine residues were replaced with glutamine (R→Q variants), appeared similar to wild type (Fig. S4). Together, these findings suggest that introduction of lysine residues into the lysine deserts leads to increased ubiquitylation.

### Ubiquitylation of introduced lysine residues is dependent on the lysine position

To test if the increased ubiquitylation is dependent on the position of the introduced lysines, we repeated the experiments with RAD23A, UBQLN1 and BAG6 variants carrying only single R→K substitutions. As expected, the effects were less dramatic. Some of the single mutant variants did not seem to affect ubiquitylation compared to the wild-type protein. However, at other positions, e.g. RAD23A R275K, UBQLN1 R177K and BAG6 R719K, increased ubiquitylation was observed (Fig. 4A). In parallel, we analyzed both the wild type and the variants carrying multiple R→K substitutions in RAD23A and UBQLN1 by mass spectrometry. We observed that one and four of the naturally occurring lysine residues for RAD23A and UBQLN1, respectively, were ubiquitylated. In all cases, these are located within the UBL domain of both proteins (Fig. 4BC) (Fig. S5). In the case of the R→K variants, we identified two additional ubiquitylation sites at the introduced lysine residues. For RAD23A, these were on K185 and K193, both located in the first UBA domain. UBQLN1 was ubiquitylated on K236 and K257, which are located in disordered regions between the structured domains (Fig. 4BC). Together with the precipitation and Western blotting results (Fig. 4A), these findings suggest that some, but importantly not all, of the artificially introduced lysine residues are susceptible to ubiquitylation. Therefore, it appears that both the position and the number of substitutions to lysine are important for the extent of ubiquitylation.

**Fig. 4.**
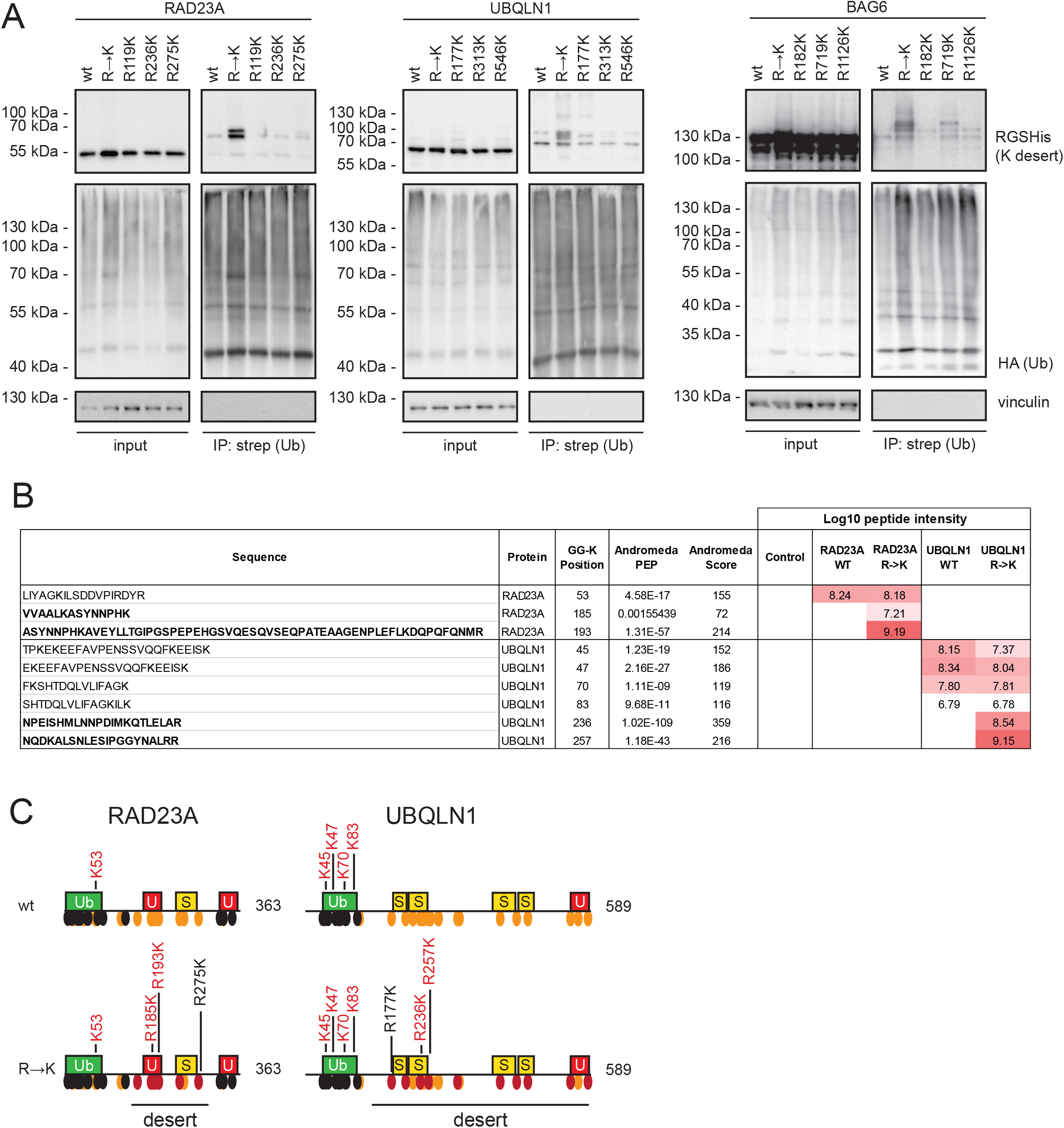
Ubiquitylation is dependent on the position of the introduced lysine. (A) U2OS cells were transiently co-transfected with HA-strep-ubiquitin and the indicated constructs. After 24 h, cells were treated with 10 μM BZ for 16 h and then used for denaturing immunoprecipitation using StrepTactin beads, followed by Western blotting. Vinculin served as a loading control. (B) The wild-type and R→K versions of Myc-tagged RAD23A and UBQLN1 were immunoprecipitated with Myc-trap beads, and peptides were analyzed by mass spectrometry for ubiquitylation. The Log_10_ peptide intensities of the identified RAD23A and UBQLN1 peptides are listed. The ubiquitylation sites in the lysines deserts are marked in bold. (C) Schematic illustration of RAD23A and UBQLN1, indicating the positions of modified lysine residues identified by mass spectrometry (red lettering) and by Western blotting (black lettering). Domain organizations as in Fig. 3A.

### The E6AP and RNF126 E3s mediate ubiquitylation of lysines introduced in lysine deserts

Next, we sought to identify E3 enzymes that are involved in catalyzing the ubiquitylation of the UBL domain substrate shuttles. Since human RAD23A was previously shown to be ubiquitylated by the E3 E6AP^33^ (UBE3A), we tested if ubiquitylation of the lysine version of RAD23A was affected upon overexpression of E6AP. Indeed, RAD23A ubiquitylation was increased when E6AP was overexpressed, which was especially pronounced for the lysine version (Fig. 5A). This effect was specific for active E6AP since we did not detect any effect on RAD23A ubiquitylation when a catalytically inactive variant (C843A) of E6AP was used (Fig. 5B). This suggests that the lysine desert in RAD23A may have evolved to protect the protein from spurious ubiquitylation by proximal E6AP and possibly other E3s.

**Fig. 5.**
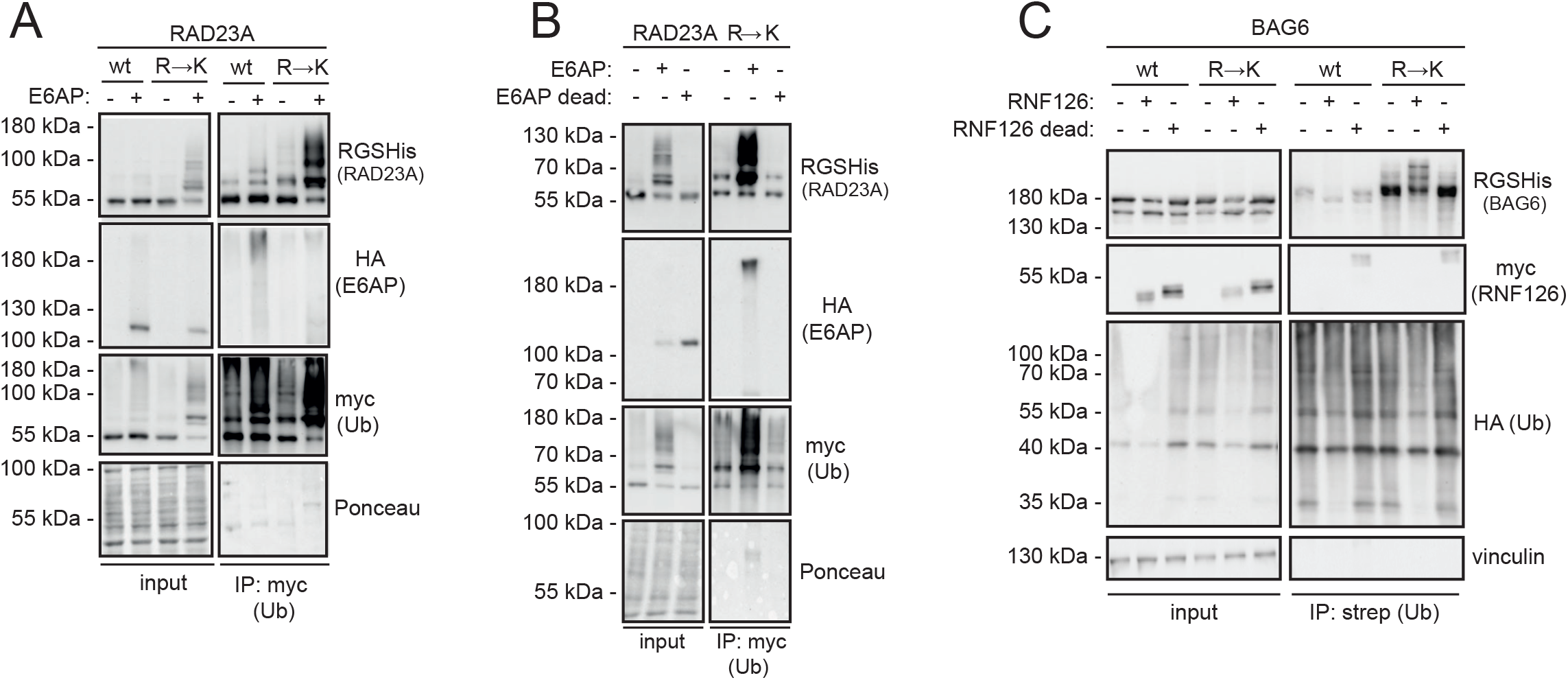
Distinct E3s catalyze the ubiquitylation of RAD23A and BAG6. (A) U2OS cells were transiently co-transfected with strep-Myc-ubiquitin, either wild-type RAD23A or the R→K variant and wild-type E6AP or empty vector. After 2 days, cells were harvested and used for denaturing immunoprecipitation with Myc-trap beads, followed by Western blotting. Staining of the membrane with Ponceau S served as a loading control. (B) U2OS cells were transiently co-transfected with strep-Myc-ubiquitin, the RAD23A R→K variant and either wild-type E6AP, a catalytically inactive (C843A) E6AP mutant or empty vector. After 2 days, cells were harvested and used for denaturing immunoprecipitation with Myc-trap beads, followed by Western blotting. Staining of the membrane with Ponceau S served as a loading control. (C) U2OS cells were transiently co-transfected with HA-strep-ubiquitin, either wild-type BAG6 or the R→K variant and with either wild-type RNF126, a catalytically inactive (C229/232A) RNF126 mutant or the empty vector. After 24 h, cells were treated with 10 μM BZ for 16 h and then used for denaturing immunoprecipitation with StrepTactin resin, followed by Western blotting. Vinculin served as a loading control.

Since human BAG6 has been reported to interact with the E3 enzyme RNF126 via its UBL domain^34^, we tested if RNF126 overexpression would affect BAG6 ubiquitylation. As a control, we included a catalytically inactive variant (C229/232A) of RNF126. Overexpression of wild-type RNF126, but not catalytically inactive RNF126, led to increased ubiquitylation of specifically the lysine version of BAG6 (Fig. 5C). We conclude that RNF126 is capable of catalyzing ubiquitylation of BAG6, and BAG6 may therefore have evolved the lysine desert to avoid RNF126 catalyzed ubiquitylation.

### Introduction of lysine residues into lysine deserts leads to an increased proteasomal degradation of UBQLN1 and BAG6, but not of RAD23A

The best-characterized role of ubiquitin is to target proteins for degradation by the proteasome. Since we noted that the introduction of lysine residues into the lysine deserts of UPS components leads to increased ubiquitylation, it is possible that this in turn leads to degradation. For more accurate determination of steady-state protein levels, we used site-specific genomic integration^35,36^ to generate stable HEK293T cell lines expressing C-terminally GFP-tagged wild-type or mutant RAD23A, UBQLN1 or BAG6. To correct for cell-to-cell variations in expression, the expression constructs also expressed mCherry from an internal ribosomal entry site (IRES) placed downstream of the GFP fusions (Fig. 6A).

**Fig. 6.**
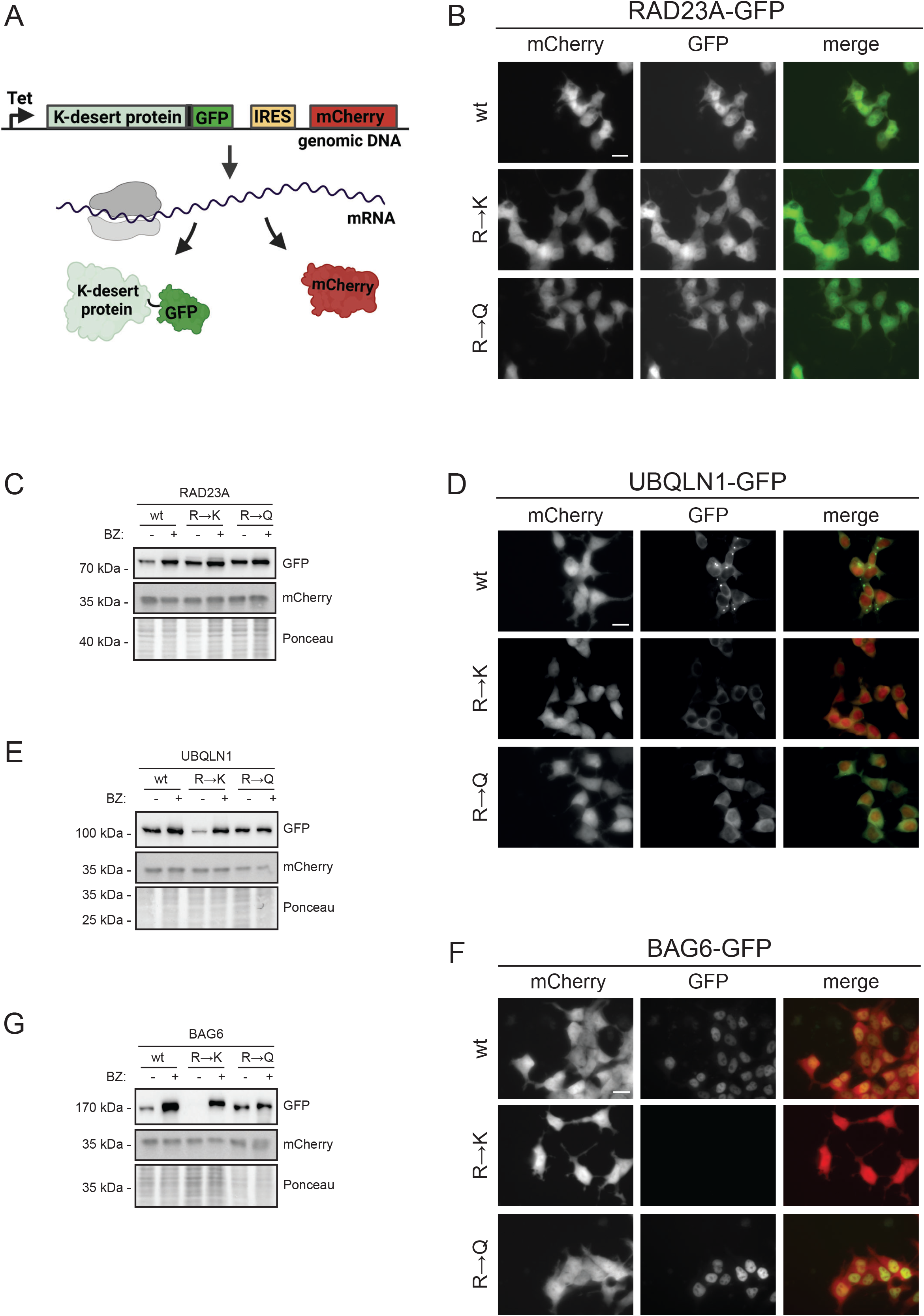
The UBQLN1 and BAG6 lysine deserts protect the proteins from proteasomal degradation. (A) The employed expression system relies on Bxb1-catalyzed site-specific plasmid integration at a “landing pad” in the genome of HEK293T cells^36^. Once the plasmid is integrated, the lysine desert protein (K-desert protein) is produced from a doxycyclin–regulated promoter (Tet) as a GFP fusion. To identify transfected cells and to correct for cell-to-cell variations in expression, the constructs also produced mCherry from an internal ribosomal entry site (IRES) located downstream of the GFP-fusions. This panel was generated using BioRender. (B) Representative fluorescence micrographs of human HEK293T cells after integration of the indicated expression constructs for RAD23A wild-type (wt), R→K, and R→Q fused to GFP. Scale bar, 20 μm. (C) Western blotting of whole cells lysates of HEK293T cells stably transfected to express the indicated expression constructs for RAD23A wild-type (wt), R→K, and R→Q fused to GFP. The blots were probed with antibodies to GFP (RAD23A) and as a control with antibodies to mCherry. Ponceau S staining of the membrane served as a loading control. (D) As panel (B), but with UBQLN1 expression. (E) As panel (C), but with UBQLN1 expression. (F) As panel (B), but with BAG6 expression. (G) As panel (C), but with BAG6 expression.

By fluorescence microscopy (Fig. 6B) and western blotting (Fig. 6C), the levels of RAD23A appeared largely unaffected by substituting arginine residues to either lysine (R→K) or glutamine (R→Q). This indicates that in case of RAD23A, ubiquitylation of the lysine residues in the lysine desert does not lead to proteasomal degradation. However, for UBQLN1 (Fig. 6DE) and BAG6 (Fig. 6FG), the R→K variants, but not the R→Q variants, were less abundant and stabilized by the proteasome inhibitor bortezomib (BZ). Hence, for these proteins, ubiquitylation of lysine residues inserted in the lysine deserts leads to proteasomal degradation of the proteins. For BAG6 we noted that the level of wild-type protein was also increased upon treating with bortezomib (Fig. 6G), indicating that BAG6 is naturally unstable but further destabilized by the introduction of lysine residues.

By fluorescence microscopy, the subcellular localization of RAD23A appeared unaffected by substituting arginines for lysine or glutamine (Fig. 6B). However, the UBQLN1 R→Q variant, although stable (Fig. 6E), appeared evenly distributed in the cytosol (Fig. 6D), whereas wild-type UBQLN1 formed cytosolic speckles (Fig. 6D), which may indicate condensate formation^37^ similar to that reported for UBQLN2^38^. BAG6 localized primarily to the nucleus, and its localization was not affected by the R→Q substitutions (Fig. 6F).

In conclusion, these data suggest that the lysine deserts protect UBQLN1 and BAG6 from proteasomal degradation, while the lysine desert in RAD23A does not appear to share this function. Moreover, the arginines in the UBQLN1 lysine desert are critical for UBQLN1 localization in cytosolic speckles, in line with the observation that arginines may help drive formation of cellular condensates^39^.

### The RAD23A lysine desert is required for optimal function in *S. pombe*

Since the lysine desert in RAD23A seems to protect the protein from ubiquitylation but does not affect its cellular stability, we reasoned that the ubiquitylation of the lysine desert might lead to impaired protein function. To test this prediction, we used the fission yeast *Schizosaccharomyces pombe*, where the deletion of the *RAD23A* orthologue, *rhp23*, results in a UV-sensitive phenotype^40–45^.

RAD23A and Rhp23 are conserved and both possess an extended lysine desert stretching between the UBL domain and the C-terminal UBA domain (Fig. S6A). When overexpressing RAD23A or its K and Q variants in *rhp23*Δ cells, we observed that the K variant, specifically, appeared to be modified in western blots of whole-cell lysates (Fig. 7A). As expected, the *rhp23*Δ strain exhibits an enhanced sensitivity towards UV radiation, which was complemented by both the RAD23A wild-type and the R→Q variant (Fig. 7B). However, in comparison, the RAD23A R→K cells appeared more sensitive to UV irradiation than the cells carrying the wild-type or R→Q variant, suggesting that the RAD23A R→K variant is partially compromised in function. In conclusion, the RAD23A lysine desert appears to be required to avoid ubiquitylation and for optimal RAD23A function in DNA repair, but is not required to protect RAD23A from proteasomal degradation.

**Fig. 7.**
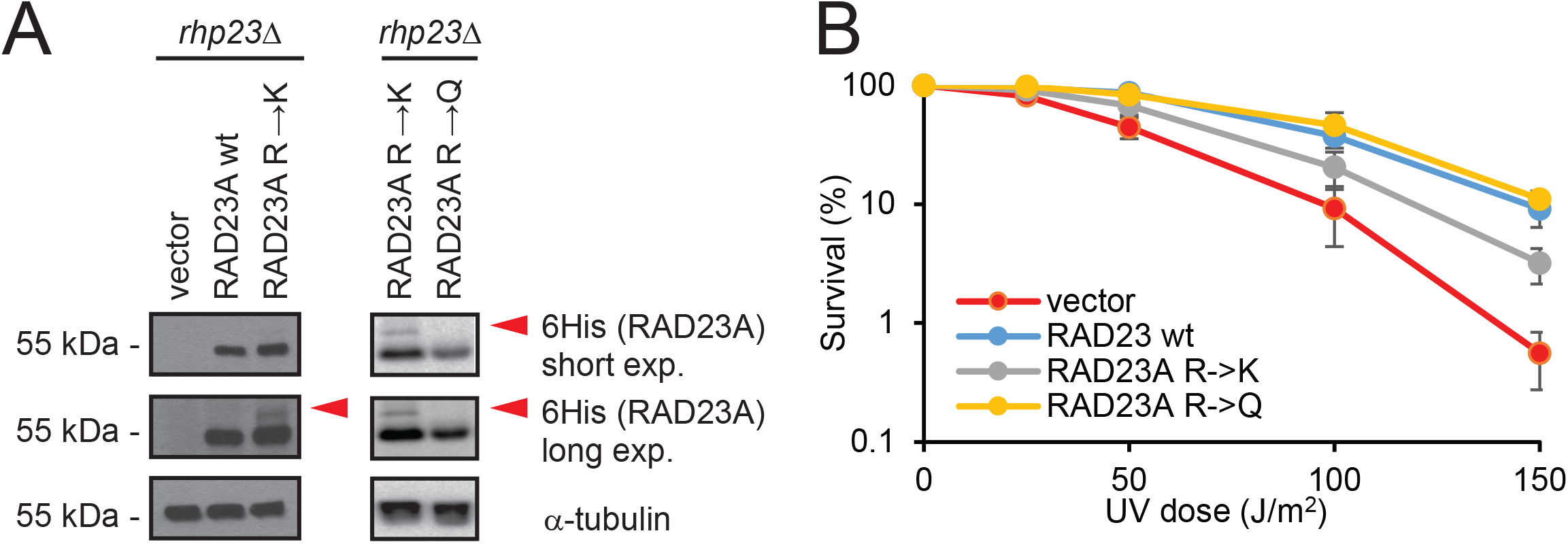
The RAD23A lysine desert is required for optimal function. (A) Western blot of RAD23A protein levels in an *rhp23*Δ *S. pombe* strain transformed with the indicated RAD23A fusion constructs. Blotting for α-tubulin served as a loading control. (B) The survival after UV radiation of exponential phase *rhp23*Δ yeast cells transformed with the indicated constructs. The error bars indicate the standard deviation (n=3).

## Discussion

Lysine deserts constitute regions in many hundreds of human proteins and may have evolved for various functions, including the evasion of lysine-linked PTMs or to maintain a specific protein structure. In this study, however, we describe the prevalence of lysine deserts as a conserved evolutionary mechanism to evade adventitious ubiquitylation. We focus on UPS components, reasoning that they are likely to become inadvertently modified, which might be particularly relevant for IDRs, where a lysine residue would be easily accessible. Accordingly, we note that it was recently reported that lysine deserts are also widespread in prokaryotes that harbor the ubiquitin-related pupylation system^46^.

Even though lysine exhibits a relatively high propensity for α-helices^47,48^, it is rated as a disorder-promoting amino acid, and IDRs have been shown to be enriched for lysines^27^. The fact that lysine is otherwise frequent in IDRs makes the existence of lysine deserts in IDRs even more remarkable and may represent a favorable trade-off, especially in the case of certain E3 enzymes. As shown for San1^17^, the IDRs in this enzyme allows for high conformational flexibility and direct substrate interaction, while the lysine deserts in the IDRs prevent its auto-ubiquitylation and degradation. We observed increased ubiquitylation when lysine was introduced into lysine deserts, but in the case of RAD23A, this did not lead to enhanced degradation. A possible reason, in agreement with previous observations on this protein^49,50^, might be that RAD23A can escape proteasomal degradation since it lacks accessible disordered regions at its termini which are needed for the proteasome to engage its substrates. Hence, the lysine desert in RAD23A appears not to be essential to protect from degradation but is still necessary for function. One possible explanation may be that ubiquitylation of lysine residues in the UBL domain of RAD23A seems to be required for function, but without affecting its degradation^51,52^. Modification at other lysine residues on the other hand may interfere with downstream signaling or act as a competitive acceptor for incoming ubiquitin. Thus, in some cases, perhaps lysine deserts have evolved to ensure focused ubiquitylation of a target protein at specific positions.

Although the lysine desert proteins contain long stretches without lysine, the position of an introduced lysine residue still appears to be critical. This dependence on position is likely a consequence of how exposed the particular position is and its proximity to the ubiquitylation machinery. Though we find that the lysine deserts tend to overlap with disordered regions, importantly even such areas of a protein may still form stable structures when interacting with binding partners. Finally, it is also possible that ubiquitylation occurs on any introduced lysine residue, albeit for some positions at a level below the detection limit.

We found that overexpression of the E3 E6AP leads to increased ubiquitylation of RAD23A, especially when lysines were introduced into the lysine desert. Although no E6AP orthologue is found in yeast, we found that RAD23A is still modified when expressed in *S. pombe*, showing that at least one additional E3 may target the RAD23A desert. As the budding yeast homologs of the two UBL-UBA shuttles, RAD23A and UBQLN1 (Rad23 and Dsk2) were shown to bind via their UBL domains to the E3 ubiquitin ligase Ufd2^53^, we speculate that perhaps Ufd2 and its human orthologue UBE4B, also contribute to the observed ubiquitylation. For BAG6 we identify its binding partner, RNF126, as an E3 which can ubiquitylate lysines introduced into its lysine desert. This indicates that RNF126 must be relatively broad in its substrate selection, and accordingly we note that RNF126 itself contains a lysine desert in its disordered N-terminal region.

Aside from PTMs and despite their similar overall charge, arginine and lysine have distinct properties^54^, and arginine and lysine residues have for example been reported to display dramatically different contributions to phase separation of disordered proteins^55^. Hence, although both lysine-rich and arginine-rich sequences can form condensates, arginine to lysine substitutions have been reported to result in a decreased propensity to phase separate^55–58^ and arginine-rich condensates are more viscous^59^. UBQLN2, and likely its paralogues, form condensates of functional importance^38,60,61^. Accordingly, we observed that overexpressed GFP-tagged UBQLN1 localizes in cytosolic speckles. The arginine residues in the UBQLN1 lysine desert appear to be critical for speckle formation since this ability is lost upon substitutions to glutamine. Though we did not observe speckle formation for the UBQLN1 lysine variant, this difference in localization may be, due to the increased degradation, leading to the UBQLN1 abundance being below a critical threshold for phase separation. Additionally, the lysine residues may alter the viscoelastic properties of the condensates, impacting the kinetics of their formation.

While we focus on the prominent group of UPS lysine deserts, we noticed that, in general, most proteins harboring pronounced lysine deserts were extracellular or plasma membrane proteins. Since the UPS is exclusively cytosolic and nuclear, there is likely a different underlying mechanism explaining why nature selected against lysine in these extracellular proteins. However, even for this group it is still tempting to speculate that the avoidance of lysine-linked modifications is a major driving force. For instance, among these proteins we noticed the extracellular lysyl oxidase LOXL1, which catalyzes the cross-linking of extracellular matrix proteins via substrate lysine residues^62^. LOXL1 possesses an extended lysine desert stretching the first two-thirds of the protein, which also overlaps with a predicted IDR. It is possible that the LOXL1 lysine desert has evolved to prevent itself from cross-linking non-specifically to extracellular matrix (ECM) components, which would likely limit its diffusion in the ECM.

The current work has generated a long list of lysine desert proteins, whose functional roles remain to be deciphered. Hopefully, future studies on these proteins will reveal more about the function of the lysine depleted regions. Along this line of thought, it would be interesting to investigate if also other types of deserts, e.g. serine or threonine deserts exist, and if so, if these have evolved to avoid undue modifications targeting these residues. Moreover, we note that several of the UPS lysine deserts, including RAD23A and UQBLN1, are also depleted for cysteine (Fig. S6AB). As cysteine is generally rare in IDRs^31^, this is likely connected with the disordered nature of the lysine deserts. Next to the observations that some proteins are targeted for ubiquitin-independent proteasomal degradation^63^ there are, however, several reports showing that ubiquitylation can also occur on the N-terminus and on serine, threonine or cysteine residues.^64–72^. Recently, Szulc *et al*. reported non-lysine ubiquitylation of the lysine desert E3s VHL and SOCS^46^, suggesting that some lysine deserts may have evolved as a means to ensure non-lysine ubiquitylation. The concomitant depletion of cysteine residues might potentially provide an evolutionary mechanism to avoid ubiquitylation of cysteine residues or other types of cysteine modifications, including oxidation.

## Materials and methods

### Bioinformatics

Lysine deserts were identified using a simple regular expression ([^K]+) on proteome files downloaded from UniProt^73^ (human: UP000005640_9606, retrieved 01/18/2021; yeast: UP000002311_559292, retrieved 01/04/2022). The entropy was calculated by considering the residues in a sequence as independent observations from an underlying categorical distribution (20 outcomes) and reported in bits. The filtering was based on a Shannon entropy cut-off of 3.7. Disorder annotations for the proteomes were extracted from the MobiDB database^28^ (retrieved at 2021/08/26 and 2022/01/04 for human and yeast, respectively). We used the “prediction-disorder-th_50” annotation, which marks residues as disordered if at least 50% of the underlying annotations are disordered for that position. Orthologues were extracted using Ensembl^74^, filtering for a specific set of diverse species, and aligned using the muscle multiple sequence alignment algorithm ^75^. Domain annotations were retrieved from the SMART database^76^. Gene ontology values were retrieved from UniProt using the Python BioServices module (v 1.7.11)^77^. UPS components were selected based on the following GO terms: Gene ontology (biological process) GO:0010498 (proteasomal protein catabolic process), GO:0006511 (ubiquitin-dependent protein catabolic process), GO:0016567 (protein ubiquitination) and/or gene ontology (molecular function) GO:0031593 (polyubiquitin modification-dependent protein binding), GO:0070628 (proteasome binding), GO:0031625 (ubiquitin protein ligase binding), GO:1990381 (ubiquitin-specific protease binding), GO:0061630 (ubiquitin protein ligase activity), GO:0004842 (ubiquitin-protein transferase activity), GO:0034450 (ubiquitin-ubiquitin ligase activity), GO:0043130 (ubiquitin binding), GO:0031624 (ubiquitin conjugating enzyme binding), GO:1990756 (ubiquitin ligase-substrate adaptor activity). The disorder predictions presented in Figure 3 were from IUPred2A^78^.

Evaluation of evolutionary conservation levels were performed measuring evolutionary distance of single site variants from the wild-type sequences of RAD23A (UniProt ID: P54725), UBQLN1 (UniProt ID Q9UMX0) and BAG6 (UniProt ID P46379). We started with the generation of multiple sequence alignments (MSAs) with the HHblits suite^79,80^, using standard parameters and an E-value threshold of 10^−20^. The obtained MSAs were comprised of 1059 sequences for RAD23A, 1270 sequences for UBQLN1, and 470 sequences for BAG6. Two additional filters were applied to the MSAs: First, all the amino acid columns in the MSA not included in the query wild-type sequences were removed. Then all the sequences with more than 50% of gaps were removed. After these filters, 1040 (RAD23A), 623 (UBQLN1), and 169 (BAG6) sequences were included in the alignments. To generate conservation scores for all the 19 possible substitutions at each position, we used the GEMME package^29^ with standard parameters. The same procedure was applied to evaluate conservation level on ten non-desert proteins reported in Supplemental file 1. For each protein we extracted disordered regions using MobiDB “prediction-disorder-th_50” and evaluated average conservation scores from each amino acid both to lysine and to arginine, excluding from the average wild-type lysine and arginine positions and their first neighbors. The non-desert human proteins used were: HMGB3 (UniProt ID O15347), H1-O (UniProt ID: P07305), H1-2 (UniProt ID: P16403), SUB1 (UniProt ID: P53999), BASP1 (UniProt ID: P80723), C17orf64 (UniProt ID: Q86WR6), ERMN (UniProt ID: Q8TAM6), NHLH2 (UniProt ID: Q02577), YF016 (UniProt ID: A6NL46), KNOP1 (UniProt ID: Q1ED39). Distribution plots of those two groups were generated using raincloud library^81^ for Python3.

### Plasmids

The expression plasmids used are listed in the supplemental material (Table S1).

### Cell culture and transfection

U2OS cells (ATCC) and HEK293T landing pad cells^36^ were propagated in Dulbecco’s Modified Eagle medium (DMEM), supplemented with 10% bovine serum (Sigma), 5000 IU/mL penicillin, 5 mg/mL streptomycin, 2 mM glutamine and, in the case of HEK293T landing pad cells, 2 μg/mL doxycycline (Dox) (Sigma-Aldrich, D9891), in a humidified atmosphere containing 5% CO_2_ at 37 °C. Transient transfections were performed with FugeneHD (Promega) in reduced serum medium OptiMEM (Gibco). To generate stable transfectants in the HEK293T landing pad cells, 10^6^ cells in 1 mL DMEM without doxycycline were seeded into 12-well plates. On the next day, the cells were transfected using 0.1 μg of the integrase vector pNLS-Bxb1-recombinase, mixed with 0.4 μg of the pVAMP recombination plasmid, 40 μL OptiMEM and 1.6 μL FugeneHD. Two days after transfection, the cells were treated with 10 nM of AP1903 (MedChemExpress) and 2 μg/mL doxycycline for two days to select for recombinant cells.

### Denaturing immunoprecipitation

Using 10-cm dishes, transiently transfected U2OS cells co-expressing either HA-, HA-Strep- or Strep-Myc-ubiquitin along with the different target constructs (ratio of 1:1) were treated with 10 μM Bortezomib (LC Laboratories) in serum-free DMEM 24 h after transfection and 16 h prior to the harvest. The cells were washed once with PBS and harvested in 300 μL lysis buffer A (30 mM Tris/HCl pH 8.1, 100 mM NaCl, 5 mM EDTA and freshly added 0.2 mM PMSF). On ice, the samples were sonicated three times for 10 sec. Subsequently, 75 μL 8% SDS was added and the samples were boiled for 10 min with intermittent vortexing. Then 1125 μL of lysis buffer A, supplemented with 2.5% Triton X-100, was added and samples incubated on ice for 30 min. Then, the samples were centrifuged at 16000 g for 60 min at 4 °C. To prepare the agarose beads, 20 μL of 50% slurry of either anti-HA resin (Sigma), StrepTactin (Qiagen) or Myc-Trap (ChromoTek) beads were washed twice in lysis buffer A. After centrifugation, 30 μL cell extract was mixed with 12.5 μL SDS sample buffer (94 mM Tris/HCl pH 6.8, 3% SDS, 40% glycerol, 0.0075% bromophenol blue, 0.0075% pyronin G) for input. The remaining extract was transferred to the tubes containing the washed beads. The samples were tumbled overnight at 4 °C and washed four times in 500 μL lysis buffer A supplemented with 1% Triton X-100 and once with lysis buffer A (without detergent). After this last washing step, as much liquid as possible was removed and 40 μL SDS sample buffer finally added to the beads. Both input and IP samples were boiled, and analyzed by SDS-PAGE and Western blotting.

### Electrophoresis and blotting

SDS-PAGE was performed with either 8% or 12.5% acrylamide gels. For the subsequent Western blotting, 0.2 μm nitrocellulose membranes (Advantec, Toyo Roshi Kaisha Ltd.) were used. The membranes were blocked in PBS containing 5% fat-free milk powder, 0.1 % Tween-20 and 5 mM NaN_3_ for at least 30 min. Incubation with the primary antibodies, all diluted in PBS containing 5% BSA and 0.1% Tween-20, was performed overnight at 4 °C. Primary antibodies used for this study were: anti-HA (Roche, 1:3000, 11867423001), anti-RGS-His (Qiagen, 1:2000, 34610), anti-Myc (ChromoTek, 1:1000, AB_2631398), anti-Vinculin (Sigma, 1:2000, V9264), anti-α-tubulin (Merck, 1:1000, TAT-1 00020911), anti-GFP (ChromoTek, 1:1000, 3H9), anti-RFP (ChromoTek, 1:1000, 6G6). The secondary antibodies were: HRP-anti-mouse IgG (Dako, 1:5000, P0260), HRP-anti-rat IgG (Thermo Fisher Scientific, 1:5000, 31470). The blots were developed using Amersham ECL detection reagent (GE Healthcare) and a ChemiDoc Imaging System (BioRad).

### Identification of ubiquitylation sites in RAD23A and UBQLN1 by mass spectrometry

Both wild-type and the lysine variants of Myc-RGS6xHis-tagged RAD23A and UBQLN1, were purified from three confluent 10-cm dishes of transfected U2OS cells following the above protocol for denaturing immunoprecipitation using Myc-trap beads (ChromoTek). A control purification from cells transfected with empty vector alone was performed in parallel. The protein was eluted from the beads with 20 μL NuPAGE LDS sample buffer (Invitrogen) and separated by SDS-PAGE using Novex 4-20% Tris-Glycine gels (Invitrogen). The gels were stained with Coomassie Brilliant Blue (Sigma). Proteins from the regions of the gel relating to modified and unmodified forms were in-gel trypsin digested^82^, alkylated with chloroacetamide and resultant peptides were resuspended in 0.1% TFA 0.5% acetic acid before analysis by LC-MS/MS. This was performed using a Q Exactive mass spectrometer (Thermo Scientific) coupled to an EASY-nLC 1000 liquid chromatography system (Thermo Scientific), using an EASY-Spray ion source (Thermo Scientific) running a 75 μm x 500 mm EASY-Spray column at 45 ºC. Two MS runs (of 60 and 150 min) were prepared using approximately 15% total peptide sample each. A top 3 data-dependent method was applied employing a full scan (m/z 300–1800) with resolution R = 70,000 at m/z 200 (after accumulation to a target value of 1,000,000 ions with maximum injection time of 20 ms). The 3 most intense ions were fragmented by HCD and measured with a resolution of R = 70,000 (60 min run) or 35,000 (150 min run) at m/z 200 (target value of 1,000,000 ions and maximum injection time of 500 ms) and intensity threshold of 2.1×104. Peptide match was set to ‘preferred’. Ions were ignored if they had unassigned charge state 1, 8 or >8 and a 10 second (60 min run) or 25 second (150 min run) dynamic exclusion list was applied.

Data analysis used MaxQuant version 1.6.1.0^83^. Default settings were used with a few exceptions. A database of the 4 transfected proteins was used as main search space (see below) with a first search using the whole human proteome (Uniprot 73920 entries – April 2019). Digestion was set to Trypsin/P (ignoring lysines and arginines N-terminal to prolines) with a maximum of 3 missed cleavages. Match between runs was not enabled. Oxidation (M), Acetyl (Protein N-term) and GlyGly (K) were included as variable modifications, with a maximum of 4 per peptide allowed. Carbamidomethyl (C) was included as a fixed modification. Only peptides of maximum mass 8000 Da were considered. Protein and peptide level FDR was set to 1% but no FDR filtering was applied to identified sites. Manual MS/MS sequence validation was used to verify GlyGly (K) peptide identifications. Peptide intensity data were reported for each sample loaded on the original gel by pooling data from the two MS runs for only the upper (modified protein) gel slices of each lane. The protein sequences are listed in the supplemental material. The mass spectrometry proteomics data have been deposited to the ProteomeXchange Consortium via the PRIDE^84^ partner repository (available upon request to the corresponding authors).

### Fluorescence microscopy

Stable transfected HEK293T landing pad cells, simultaneously expressing C-terminally tagged GFP-constructs and mCherry from an IRES (Fig. 6A), were grown in 6-well dishes. Images of the cells were recorded directly from the 6-well plates and with a Zeiss AxioVert.A1 microscope equipped with an AxioCam ICm 1 Rev. 1 FireWire (D) camera. EGFP was excited at 475 nm and mCherry at 590 nm. Image processing was performed using ImageJ (version 1.51j8).

### Yeast strains and techniques

The fission yeast *rhp23*Δ strain^85^ was grown in standard rich media (YES media: 30 g/L glucose, 5 g/L yeast extract, 0.2 g/L adenine, 0.2 g/L uracil, 0.2 g/L leucine) at 30 °C. The cells were transformed with pREP1-based vectors using lithium acetate^86^. Transformants were cultured in Edinburgh Minimal Media 2 (EMM2) (Sunrise Science) without leucine for plasmid selection. For the survival assays, yeast cultures were grown at 30 °C to exponential phase and spread on solid media plates and irradiated. Colonies were counted after four days at 30 °C. For western blotting, proteins were extracted as whole cell lysate using trichloroacetic acid (Sigma) and glass beads as described before^87^.

## Online material

The mass spectrometry proteomics data have been deposited to the ProteomeXchange Consortium and will be forwarded upon request to the corresponding authors.

## Supplemental material

This manuscript contains supplemental material.

Supplemental figures and tables, Fig. S1-S6, Table S1, and protein sequences.

Supplemental File 1, The human proteome sorted for lysine deserts.

Supplemental File 2, The proteomes of selected model organisms sorted for lysine deserts.

Supplemental File 3, The GEMME conservation scores for the selected proteins.

## Acknowledgements

The authors thank Sofie V. Nielsen and Birthe B. Kragelund for the helpful discussions and comments on the manuscript, and the REPIN consortium for discussions on IDPs. We thank Anne-Marie Lauridsen and Søren Lindemose for excellent technical assistance. We thank Mads Gyrd-Hansen for the HA-strep ubiquitin expression construct. We acknowledge access to computing resources from the Biocomputing Core Facility at the Department of Biology, University of Copenhagen.

## Author contributions

C.K., M.G.T., W.B., N.O., and M.H.T. conducted the experiments. Data analyses by C.K., M.G.T., W.B., M.H.T., K.H., M.C. and R.H.P. Experimental design by K.L.-L., K.H., W.B., and R.H.P. K.H., and R.H.P. conceived the study. C.K., M.G.T., and R.H.P. wrote the paper.

