## Supplemental Figures and Tables for "Lysine deserts prevent adventitious ubiquitylation of ubiquitin-proteasome components"

### *Supplemental material*


| | |
|---|---|
| *Supplemental figure, Fig. S1* | *p.2* |
| *Supplemental figure, Fig. S2* | *p.3* |
| *Supplemental figure, Fig. S3* | *p.4* |
| *Supplemental figure, Fig. S4* | *p.5* |
| *Supplemental figure, Fig. S5* | *p.6* |
| *Supplemental figure, Fig. S6* | *p.7* |
| *Supplemental Table, Table S1* | *p.8* |
| *Protein sequence details for mass spectrometry* | *p.9* |

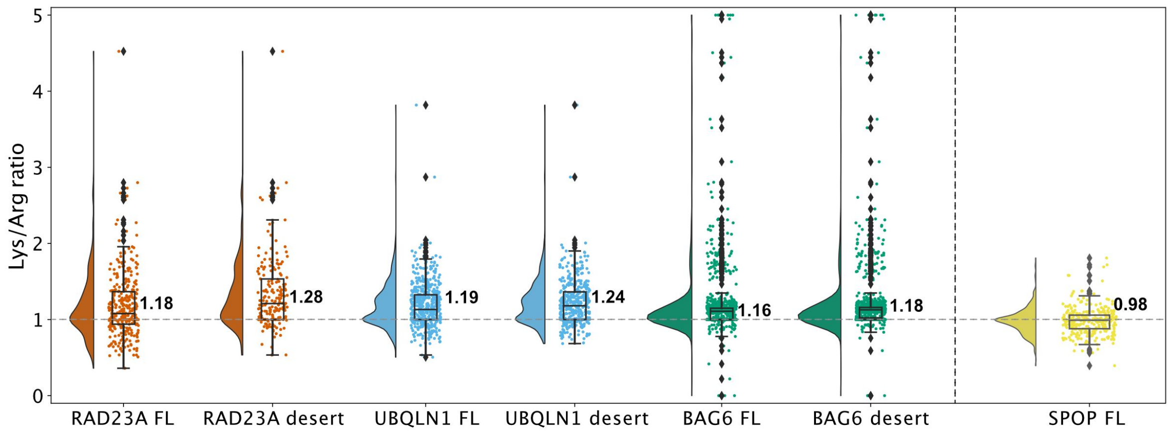


**Fig. S1.** *Sequence alignments indicate that substitutions to Lys are predicted as more unfavorable than substitutions to Arg.*

The Lys/Arg ratios for the GEMME-based conservation scores plotted for the indicated full-length (FL) proteins and lysine depleted regions (desert).

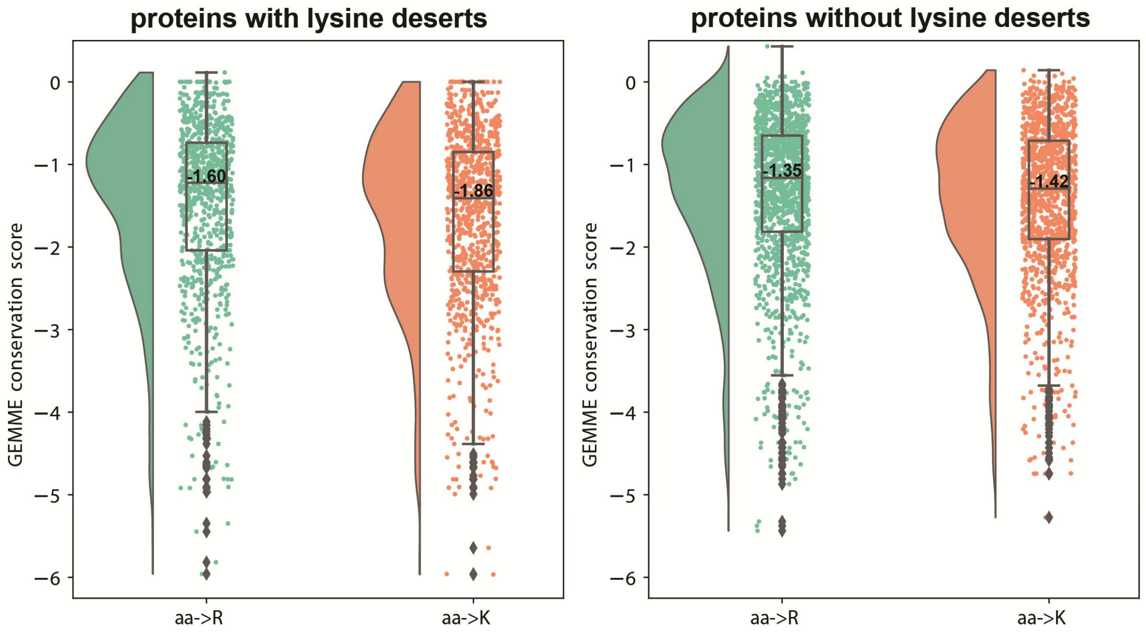


**Fig. S2.** *Introduction of lysine is more detrimental in lysine desert proteins than in non-lysine desert proteins.* GEMME conservation scores for substitutions of any amino acid in the selected proteins to R or K, presented as raincloud plots. Each plot shows the distribution of data points (left) and the raw scores (right). Additionally, in each set a boxplot reports the average, quartiles, maximum and minimum values.

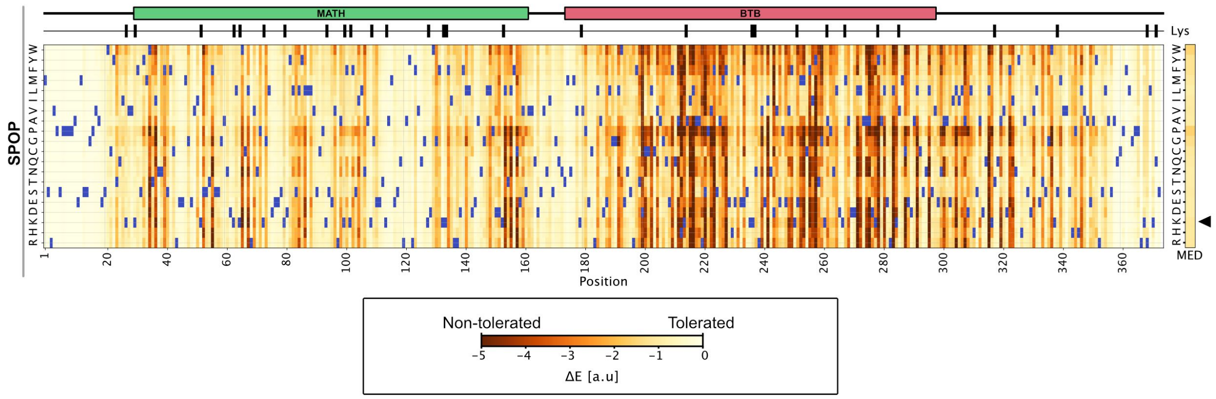


**Fig. S3.** *Example of a conservation map for a non-lysine desert protein.*
Evolutionary conservation analysis using multiple sequence alignments of the human UPS protein SPOP presented as a heatmap. GEMME scores (ΔE) close to zero (white and light yellow colors) indicate that a given amino acid substitution is compatible with the alignments, while negative scores (red and dark yellow colors) indicate that the substitution is incompatible with the alignments and therefore likely detrimental to the protein structure and/or function. The wild-type amino acid residue at each position is shown in blue. The domain organization and the positions of lysine residues are shown above. The median score (MED) for substitutions to the indicated amino acid resides across the entire protein is shown to the right. Note that substitutions to lysine (arrowhead) in general appear to be tolerated for this non-desert protein.

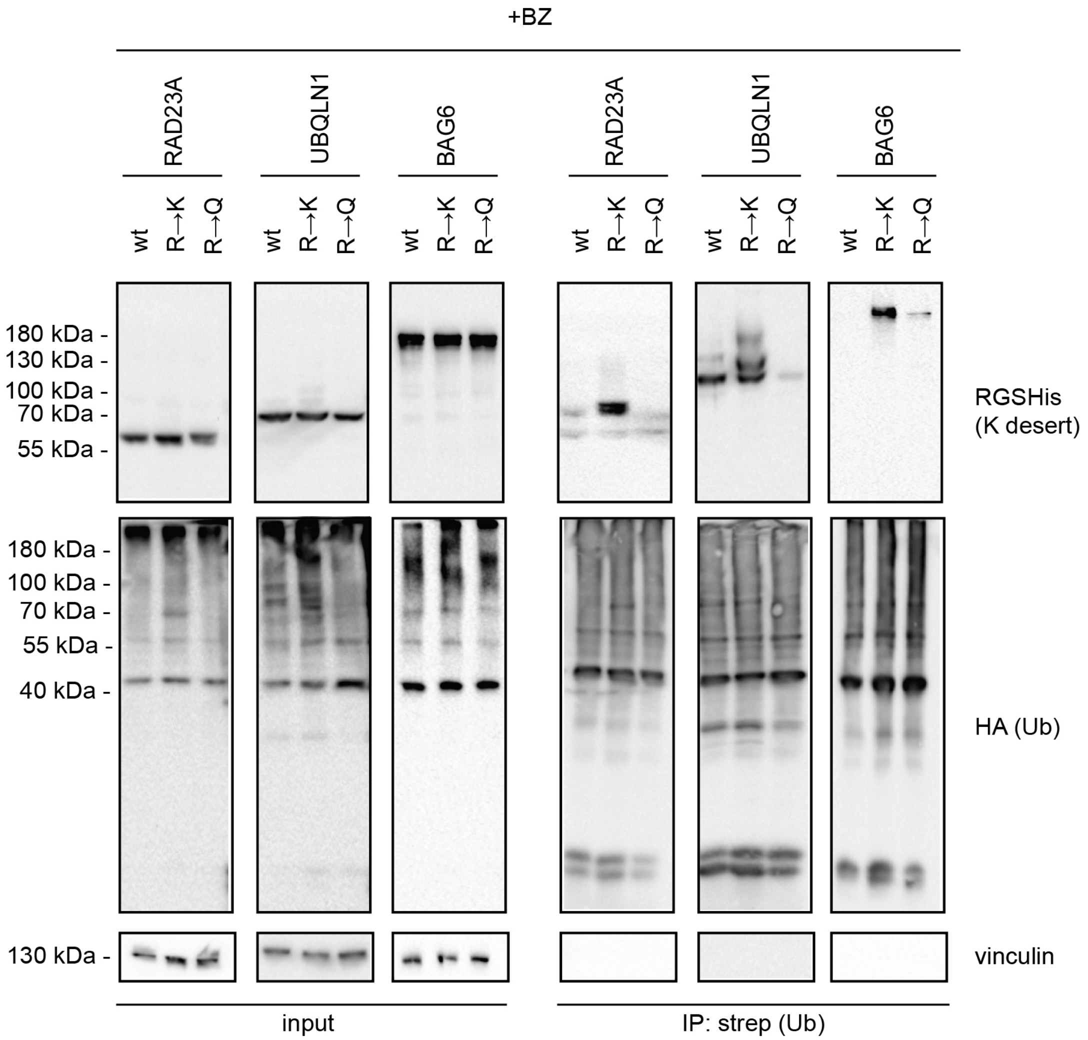


**Fig. S4.** *Introduction of lysine, but not glutamine, residues leads to ubiquitylation.*
U2OS cells were transiently co-transfected with HA-strep-tagged ubiquitin and the indicated constructs. After 24 h, cells were treated with 10 µM BZ for 16 h and then used for denaturing immunoprecipitation (IP) with StrepTactin beads. Protein was visualized by Western blotting and vinculin was used as a loading control.

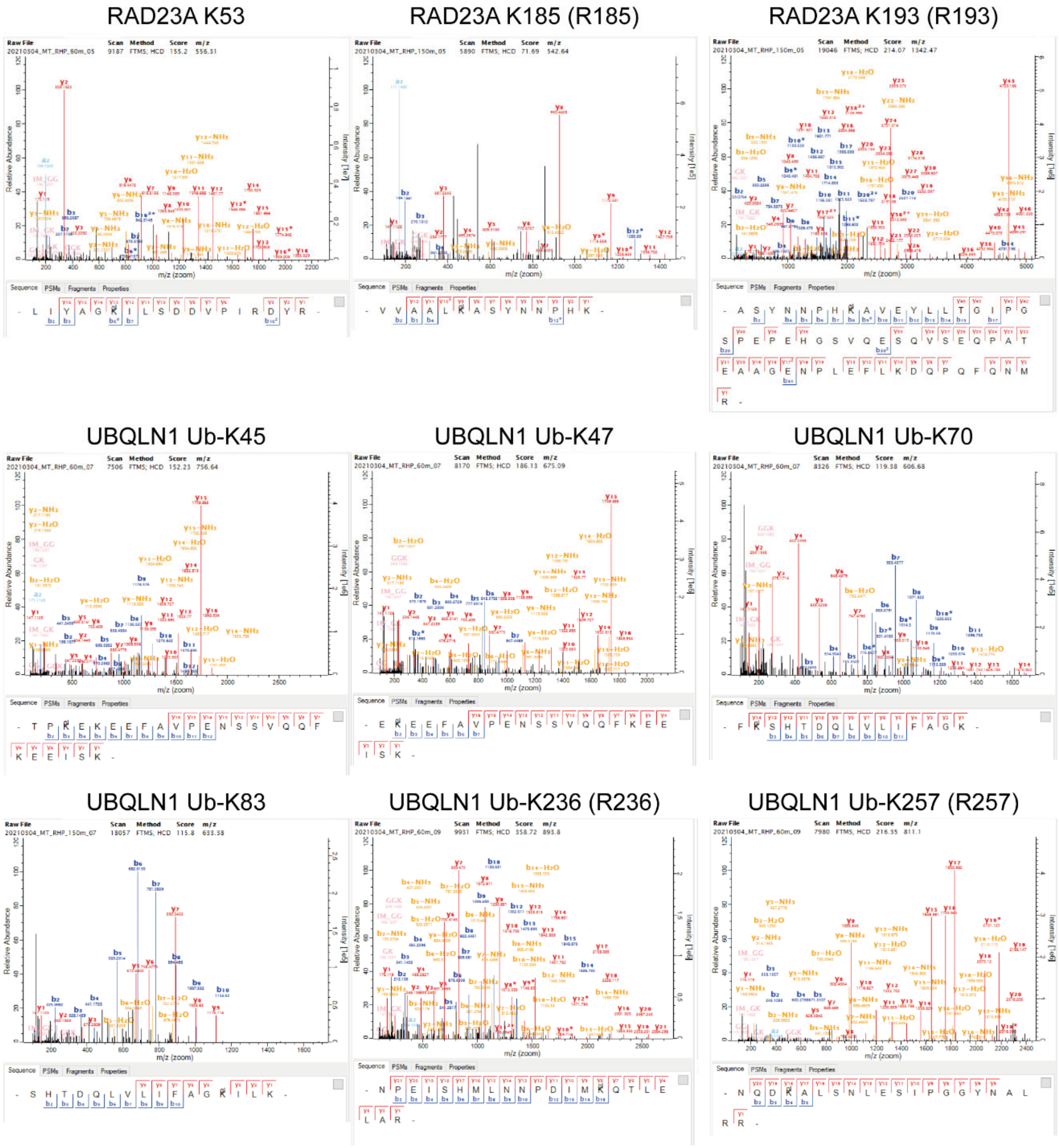


**Fig. S5.** *Annotated MS/MS spectra.*
MaxQuant annotated MS/MS spectra for data shown in Fig. 4B.

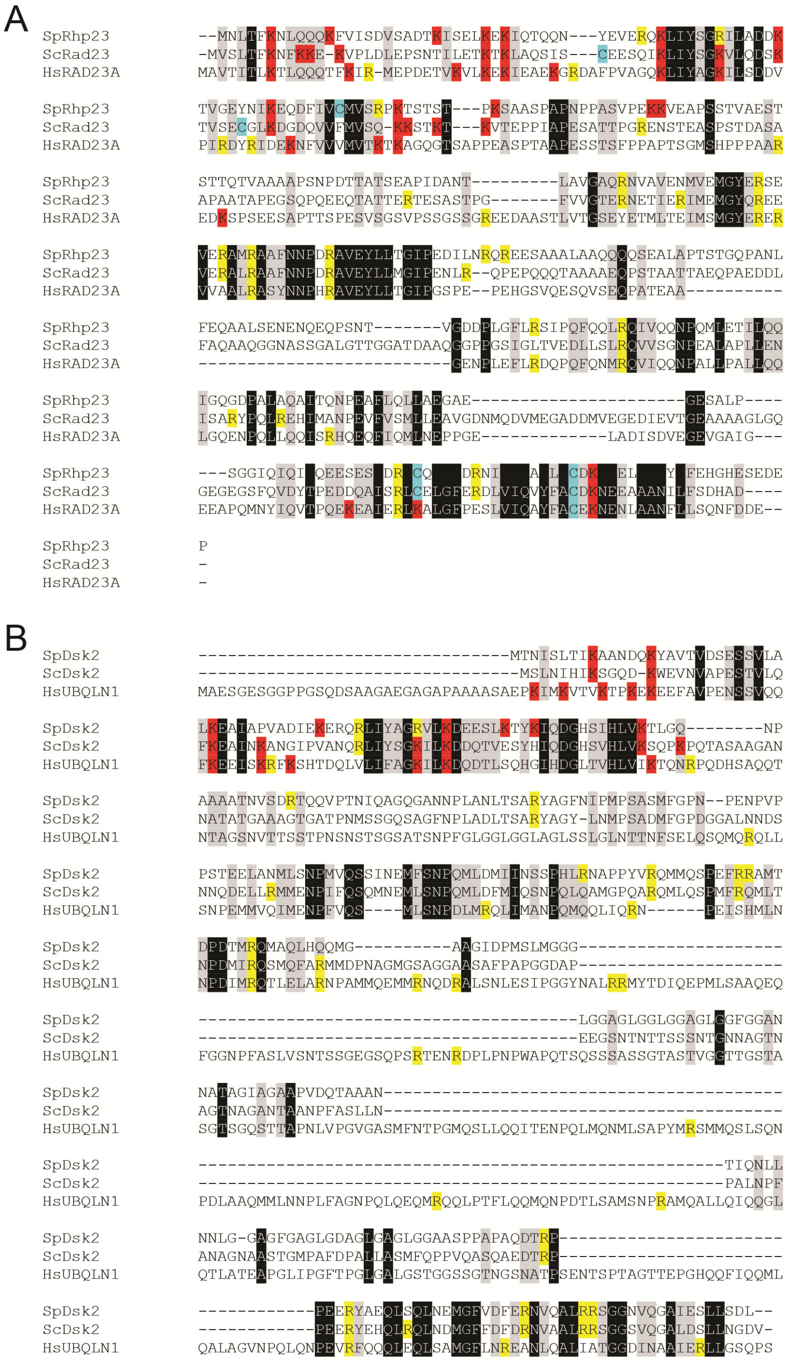

**Fig. S6.** *Phylogenetic conservation of the lysine deserts in RAD23A and UBQLN1 to yeast.*
(A) Multiple sequence alignment of human RAD23A (HsRAD23A) with its *Schizosaccharomyces pombe* (SpRhp23) and *Saccharomyces cerevisiae* (ScRad23) orthologues. Identical (black) and similar (grey) residues have been shaded. Arginine (yellow), lysine (red) and cysteine (cyan) residues have been marked. (B) Multiple sequence alignment of human UBQLN1 (HsUBQLN1) with its *S. pombe* (SpDsk2) and *S. cerevisiae* (ScDsk2) orthologues. Identical (black) and similar (grey) residues have been shaded. Arginine (yellow) and lysine (red) residues have been marked. Note that UBQLN1 and its yeast orthologues do not contain any cysteine residues. The alignments were prepared using ClustalW v.2.1.

**Table S1**

*Plasmids used in this study*

| Vector | Insert | Mutations | Tag* | Origin |
|---|---|---|---|---|
| pcDNA3.1 | RAD23A wt | none | NT-RGS6xHis | Genscript |
| pcDNA3.1 | RAD23A R→K | R149/177/179/185/193/236/275K | NT-RGS6xHis | Genscript |
| pcDNA3.1 | RAD23A R→Q | R149/177/179/185/193/236/275Q | NT-RGS6xHis | Genscript |
| pcDNA3.1 | RAD23A | R119K | NT-RGS6xHis | Genscript |
| pcDNA3.1 | RAD23A | R236K | NT-RGS6xHis | Genscript |
| pcDNA3.1 | RAD23A | R275K | NT-RGS6xHis | Genscript |
| pcDNA3.1 | RAD23A wt | none | NT-Myc-RGS6xHis | Genscript |
| pcDNA3.1 | RAD23A R→K | R149/177/179/185/193/236/275K | NT-Myc-RGS6xHis | Genscript |
| pVAMP | RAD23A wt | none | CT-GFP | Genscript |
| pVAMP | RAD23A R→K | R149/177/179/185/193/236/275K | CT-GFP | Genscript |
| pVAMP | RAD23A R→Q | R149/177/179/185/193/236/275Q | CT-GFP | Genscript |
| pcDNA3.1 | UBQLN1 wt | none | NT-RGS6xHis | Genscript |
| pcDNA3.1 | UBQLN1 R→K | R177/206/236/257/313/401/435/458/546/582K | NT-RGS6xHis | Genscript |
| pcDNA3.1 | UBQLN1 R→Q | R177/206/236/257/313/401/435/458/546/582Q | NT-RGS6xHis | Genscript |
| pcDNA3.1 | UBQLN1 | R177K | NT-RGS6xHis | Genscript |
| pcDNA3.1 | UBQLN1 | R313K | NT-RGS6xHis | Genscript |
| pcDNA3.1 | UBQLN1 | R546K | NT-RGS6xHis | Genscript |
| pcDNA3.1 | UBQLN1 wt | none | NT-Myc-RGS6xHis | Genscript |
| pcDNA3.1 | UBQLN1 R→K | R177/206/236/257/313/401/435/458/546/582K | NT-Myc-RGS6xHis | Genscript |
| pVAMP | UBQLN1 wt | none | CT-GFP | Genscript |
| pVAMP | UBQLN1 R→K | R177/206/236/257/313/401/435/458/546/582K | CT-GFP | Genscript |
| pVAMP | UBQLN1 R→Q | R177/206/236/257/313/401/435/458/546/582Q | CT-GFP | Genscript |
| pcDNA3.1 | BAG6 wt | none | NT-RGS6xHis | Genscript |
| pcDNA3.1 | BAG6 R→K | R127/174/192/242/289/393/445/513/719/780/805/901/986K | NT-RGS6xHis | Genscript |
| pcDNA3.1 | BAG6 R→Q | R127/174/192/242/289/393/445/513/719/780/805/901/986Q | NT-RGS6xHis | Genscript |
| pcDNA3.1 | BAG6 | R182K | NT-RGS6xHis | Genscript |
| pcDNA3.1 | BAG6 | R719K | NT-RGS6xHis | Genscript |
| pcDNA3.1 | BAG6 | R1126K | NT-RGS6xHis | Genscript |
| pVAMP | BAG6 wt | none | CT-GFP | Genscript |
| pVAMP | BAG6 R→K | R127/174/192/242/289/393/445/513/719/780/805/901/986K | CT-GFP | Genscript |
| pVAMP | BAG6 R→Q | R127/174/192/242/289/393/445/513/719/780/805/901/986Q | CT-GFP | Genscript |
| pcDNA5/FRT | UBQLN2 wt | none | NT-2xHA | This study |
| pcDNA5/FRT | UBQLN2 R→K | R269/617K | NT-2xHA | This study |
| pcDNA5/FRT | RNF115 wt | none | NT-2xHA | This study |
| pcDNA5/FRT | RNF115 R→K | R62/125/126K | NT-2xHA | This study |
| pcDNA5/FRT | PSMF wt | none | NT-2xHA | This study |
| pcDNA5/FRT | PSMF R→K | R219K | NT-2xHA | This study |
| pREP1 | - | none | - | Lab stock |
| pREP1 | RAD23A wt | none | NT-RGS6xHis | Genscript |
| pREP1 | RAD23A R→K | R149/177/179/185/193/236/275K | NT-RGS6xHis | Genscript |
| pREP1 | RAD23A R→Q | R149/177/179/185/193/236/275Q | NT-RGS6xHis | Genscript |
| pcDNA3.1 | Ubiquitin (Ub) | none | NT-HA-Strep | M. Gyrd-Hansen |
| pcDNA3.1 | Ubiquitin (Ub) | none | NT-Strep-Myc | Genscript |
| pcDNA3.1 | RNF126 | none | NT-Myc | Genscript |
| pcDNA3.1 | RNF126 dead | C229/232A | NT-Myc | Genscript |
| pcDNA3.1 | E6AP | none | NT-HA | TWIST |
| pcDNA3.1 | E6AP dead | C843A | NT-HA | TWIST |
| pRK5-HA | Ubiquitin (Ub) | none | NT-HA | Addgene |
| pNLS-Bxb1 | NLS-Bxb1 | - | - | D. M. Fowler |

*NT, tag at N-terminus; CT, tag at C-terminus.

**Protein sequence details for mass spectrometry**

>RAD23A_WT
MEQKLISEEDLGTATMAYYRGSHHHHHHSRSMAVTITLKTLQQQTFKIRMEPDETVKVLKEKIEAEKGRDAFP
VAGQKLIYAGKILSDDVPIRDYRIDEKNFVVVMVTKTKAGQGTSAPPEASPTAAPESSTSFPPAPTSGMSHPP
PAAREDKSPSEESAPTTSPESVSGSVPSSGSSGREEDAASTLVTGSEYETMLTEIMSMGYERERVVAALRASY
NNPHRAVEYLLTGIPGSPEPEHGSVQESQVSEQPATEAAGENPLEFLRDQPQFQNMRQVIQQNPALLPALLQQ
LGQENPQLLQQISRHQEQFIQMLNEPPGELADISDVEGEVGAIGEEAPQMNYIQVTPQEKEAIERLKALGFPE
SLVIQAYFACEKNENLAANFLLSQNFDDE

>RAD23A_RK
MEQKLISEEDLGTATMAYYRGSHHHHHHSRSMAVTITLKTLQQQTFKIRMEPDETVKVLKEKIEAEKGRDAFP
VAGQKLIYAGKILSDDVPIRDYRIDEKNFVVVMVTKTKAGQGTSAPPEASPTAAPESSTSFPPAPTSGMSHPP
PAAREDKSPSEESAPTTSPESVSGSVPSSGSSGKEEDAASTLVTGSEYETMLTEIMSMGYEKEKVVAALKASY
NNPHKAVEYLLTGIPGSPEPEHGSVQESQVSEQPATEAAGENPLEFLKDQPQFQNMRQVIQQNPALLPALLQQ
LGQENPQLLQQISKHQEQFIQMLNEPPGELADISDVEGEVGAIGEEAPQMNYIQVTPQEKEAIERLKALGFPE
SLVIQAYFACEKNENLAANFLLSQNFDDE

>UBQLN1_WT
MEQKLISEEDLGTATMAYYRGSHHHHHHSRSMAESGESGGPPGSQDSAAGAEGAGAPAAAASAEPKIMKVTVK
TPKEKEEFAVPENSSVQQFKEEISKRFKSHTDQLVLIFAGKILKDQDTLSQHGIHDGLTVHLVIKTQNRPQDH
SAQQTNTAGSNVTTSSTPNSNSTSGSATSNPFGLGGLGGLAGLSSLGLNTTNFSELQSQMQRQLLSNPEMMVQ
IMENPFVQSMLSNPDLMRQLIMANPQMQQLIQRNPEISHMLNNPDIMRQTLELARNPAMMQEMMRNQDRALSN
LESIPGGYNALRRMYTDIQEPMLSAAQEQFGGNPFASLVSNTSSGEGSQPSRTENRDPLPNPWAPQTSQSSSA
SSGTASTVGGTTGSTASGTSGQSTTAPNLVPGVGASMFNTPGMQSLLQQITENPQLMQNMLSAPYMRSMMQSL
SQNPDLAAQMMLNNPLFAGNPQLQEQMRQQLPTFLQQMQNPDTLSAMSNPRAMQALLQIQQGLQTLATEAPGL
IPGFTPGLGALGSTGGSSGTNGSNATPSENTSPTAGTTEPGHQQFIQQMLQALAGVNPQLQNPEVRFQQQLEQ
LSAMGFLNREANLQALIATGGDINAAIERLLGSQPS

>UBQLN1_RK
MEQKLISEEDLGTATMAYYRGSHHHHHHSRSMAESGESGGPPGSQDSAAGAEGAGAPAAAASAEPKIMKVTVK
TPKEKEEFAVPENSSVQQFKEEISKRFKSHTDQLVLIFAGKILKDQDTLSQHGIHDGLTVHLVIKTQNRPQDH
SAQQTNTAGSNVTTSSTPNSNSTSGSATSNPFGLGGLGGLAGLSSLGLNTTNFSELQSQMQKQLLSNPEMMVQ
IMENPFVQSMLSNPDLMKQLIMANPQMQQLIQRNPEISHMLNNPDIMKQTLELARNPAMMQEMMRNQDKALSN
LESIPGGYNALRRMYTDIQEPMLSAAQEQFGGNPFASLVSNTSSGEGSQPSKTENRDPLPNPWAPQTSQSSSA
SSGTASTVGGTTGSTASGTSGQSTTAPNLVPGVGASMFNTPGMQSLLQQITENPQLMQNMLSAPYMKSMMQSL
SQNPDLAAQMMLNNPLFAGNPQLQEQMKQQLPTFLQQMQNPDTLSAMSNPKAMQALLQIQQGLQTLATEAPGL
IPGFTPGLGALGSTGGSSGTNGSNATPSENTSPTAGTTEPGHQQFIQQMLQALAGVNPQLQNPEVKFQQQLEQ
LSAMGFLNREANLQALIATGGDINAAIEKLLGSQPS
